# Dynamic semantic cognition: Characterising coherent and controlled conceptual retrieval through time using magnetoencephalography and chronometric transcranial magnetic stimulation

**DOI:** 10.1101/168203

**Authors:** Catarina Teige, Giovanna Mollo, Rebecca Millman, Nicola Savill, Jonathan Smallwood, Piers L. Cornelissen, Elizabeth Jefferies

**Author notes:** Correspondence to: Piers Cornelissen, Psychology Department, Faculty of Health and Life Sciences, Northumbria University, Newcastle upon Tyne, NE1 8ST.

## Abstract

Distinct neural processes are thought to support the retrieval of semantic information that is (i) coherent with strongly-encoded aspects of knowledge, and (ii) non-dominant yet relevant for the current task or context. While the brain regions that support coherent and controlled patterns of semantic retrieval are relatively well-characterised, the temporal dynamics of these processes are not well-understood. This study used magnetoencephalography (MEG) and dual-pulse chronometric transcranial magnetic stimulation (cTMS) in two separate experiments to examine temporal dynamics within the temporal lobe during the retrieval of strong and weak associations. MEG results revealed a dissociation within left temporal cortex: anterior temporal lobe (ATL) showed greater oscillatory response for strong than weak associations, while posterior middle temporal gyrus (pMTG) showed the reverse pattern. In the cTMS experiment, stimulation of ATL at ~150ms disrupted the efficient retrieval of strong associations, indicating a necessary role for ATL in coherent conceptual activations. Stimulation of pMTG at the onset of the second word disrupted the retrieval of weak associations, suggesting this site may maintain information about semantic context from the first word, allowing engagement of semantic control. Together these studies provide converging evidence for a functional dissociation within the temporal lobe, across both tasks and time.

---

Semantic cognition allows us to understand the meaning of our environment to drive appropriate thoughts and behaviour. It comprises several distinct yet interacting components (Jefferies, 2013; Jefferies & Lambon Ralph, 2006; Lambon Ralph, Jefferies, Patterson & Rogers, 2017). Semantic representations capture the meanings of words and objects across contexts, supporting coherent conceptual retrieval from fragmentary inputs and generalisation across situations. However, the retrieval of specific aspects of knowledge in a context-dependent fashion requires control mechanisms that shape evolving retrieval towards currently-pertinent semantic features, and away from dominant yet irrelevant associations. While patterns of activation within the semantic store may be sufficient to uncover links between items that are highly coherent with long-term knowledge (i.e. items that share multiple features or are frequently associated, such as *pear*-*apple* or *tree*-*apple*), additional control processes may be required to recover non-dominant aspects of knowledge, such as *worm*-*apple*, since strong but currently-irrelevant associations (e.g. *worm*-*soil*) must be disregarded (Lambon Ralph, Jefferies, Patterson & Rogers, 2017; Gold et al., 2006).

Although the brain regions that support semantic cognition are relatively well-described, the temporal dynamics are not. Neuroimaging studies have highlighted the importance of a distributed left-dominant network underpinning semantic cognition, including anterior temporal lobe (ATL), posterior middle temporal gyrus (pMTG) and inferior frontal gyrus (LIFG) (Jefferies, 2013; Vandenberghe et al., 1996; Xu, Qixiang, Zaizhu, Yong, & Yanchao, 2016; Lambon Ralph et al., 2017; Binder, Desai, Graves & Conant, 2009). These brain regions make dissociable contributions to semantic cognition, although their specific roles remain controversial. The ventral ATL is proposed to support heteromodal concepts that are extracted from multiple inputs (e.g., vision, audition, smell; Patterson, Nestor & Rogers, 2007; Lambon Ralph et al., 2017). Patients with semantic dementia (SD), show progressive degradation of knowledge across modalities following atrophy and hypometabolism in ATL (Mion et al., 2010; Bozeat et al., 2000; Rogers et al., 2006). Convergent evidence for a role of ATL in multimodal conceptual processing is provided by positron emission tomography (e.g. Bright et al., 2004; Crinion et al., 2003; Devlin et al., 2000; Noppeney & Price, 2002; Rogers et al., 2006; Scott et al., 2000), functional magnetic resonance imaging (fMRI) – particularly when magnetic susceptibility artefacts within ATL are minimised (Binney et al., 2010; Murphy et al., 2017; Visser et al.,2010; 2012), magnetoencephalography (MEG) (Lau et al, 2013; Clarke et al., 2011; Marinković et al., 2003; Fujimaki et al., 2009; Mollo et al., 2017), intracranial electrode arrays (Chan et al., 2011; Chen et al., 2016) and transcranial magnetic stimulation (TMS) (Lambon Ralph et al., 2009; Pobric et al., 2009; Jackson, Lambon Ralph & Pobric, 2016). The ATL is allied to the default mode network (DMN) in terms of connectivity and function (Binder et al., 2003; Davey et al., 2015; 2016; Wirth et al., 2011), although the maximal semantic response in ATL is not identical to the site of peak DMN connectivity (Jackson et al., 2016). In common with DMN regions, ATL shows a larger response to easy or more automatic aspects of semantic retrieval, such as identifying dominant aspects of knowledge (e.g., linking DOG with CAT; Davey et al., 2016), and when coherent meaning emerges from conceptual combinations (Bemis & Pylkkänen, 2012; Hoffman, Binney & Lambon Ralph, 2015). ATL is also implicated in semantic retrieval during mind-wandering (Binder et al., 1999; Smallwood et al., 2016). Collectively, these findings suggest that ATL responds most strongly when ongoing semantic retrieval is highly coherent with long-term knowledge – although *causal* evidence is currently lacking.

Brain regions distinct from ATL are implicated in the *control* of semantic cognition. The contribution of left inferior prefrontal cortex to executive-semantic processes has been documented by many fMRI studies (e.g., Thompson-Schill, D’Esposito, Aguirre &Farah, 1997; Badre, Poldrack, Pare-Blagoev, Insler & Wagner, 2005; Noppeney, Phillips & Price, 2004; Noonan et al., 2013; Bedny, McGill & Thompson-Schill, 2008). Convergent evidence for a causal contribution of this region has been provided by transcranial magnetic stimulation (TMS, Hoffman, Jefferies & Lambon Ralph, 2010; Whitney et al., 2011) and neuropsychology: patients with damage to LIFG have difficulty flexibly tailoring their semantic retrieval to suit the circumstances (Thompson-Schill et al., 1998; Jefferies & Lambon Ralph, 2006; Corbett, Jefferies & Lambon Ralph, 2009; Thompson et al., 2015). While the contributions of ATL and LIFG align with recent component process views of semantic cognition (e.g., the Controlled Semantic Cognition framework, which suggests semantic cognition reflects an interaction of conceptual representations and control processes, Lambon Ralph et al., 2017), the contribution of pMTG remains controversial. Some accounts have proposed that posterior temporal areas provide an important store of conceptual representations (Martin, 2007), with pMTG specifically implicated in knowledge of actions and events (Chao, Haxby & Martin, 1999; Martin et al., 1995). Alternatively, a growing literature supports the view that pMTG is part of a distributed network with LIFG and other regions underpinning semantic control (Vitello et al., 2014; Jefferies, 2013; Davey et al., 2016; Noonan et al., 2013; Gold et al., 2006). A recent meta-analysis showed that a widely distributed set of cortical regions is reliably activated across diverse manipulations of semantic control demands, with pMTG showing the second most consistent response after LIFG (Noonan et al., 2013). Semantic control deficits can follow from either left prefrontal or posterior temporal lesions (Jefferies & Lambon Ralph, 2006; Noonan et al., 2010); moreover, inhibitory TMS to pMTG and LIFG produces equivalent disruption of semantic judgements that require controlled but not automatic retrieval (Whitney et al., 2011; Davey et al., 2015), and inhibitory stimulation of LIFG elicits a compensatory increase in pMTG (Hallam et al., 2017). These regions also show a strong pattern of both structural and functional connectivity (Davey et al., 2016; Jeyoung & Lambon Ralph, 2016), consistent with the view that they form a large-scale distributed network underpinning controlled aspects of semantic retrieval. This semantic control network partially overlaps with the frontoparietal control network and thus both LIFG and pMTG have a different pattern of large-scale connectivity from ATL (Davey et al., 2016).

Component process accounts of semantic cognition (e.g. Jefferies, 2013; Lambon Ralph et al., 2017) predict a functional dissociation within the temporal lobe – with ATL supporting efficient retrieval when currently-relevant semantic information is highly coherent with dominant aspects of long term knowledge, and pMTG playing a critical role at times when such knowledge cannot serve the goal of the moment. The current work tests this predicted dissociation between temporal lobe sites by examining how their contribution to semantic processing changes when dominant conceptual associations no longer support appropriate patterns of retrieval, and more weakly-encoded information is required. We presented two words successively and manipulated the strength of the relationship between them. When two words are strongly associated, retrieval of the relevant conceptual link is thought to be relatively automatic, since the meaning of the second word strongly overlaps with features activated from the first word. For weaker associations, predictions from the first word are less useful: there may be partial overlap between the meaning of the second word and conceptual features activated from the first word, but irrelevant, dominant semantic associations will have stronger activation, and thus controlled semantic retrieval might be required to focus processing on currently-relevant features (cf. Badre et al., 2005; Whitney et al., 2011). This paradigm provides a clear temporal marker (the onset of the second word) from which to examine more coherent and controlled patterns of semantic retrieval.

If the two temporal lobe regions play distinct roles in automatic and controlled semantic retrieval, we reasoned that, as well as overall differences in their response to strong and weak associations, there might also be differences in the timing of these effects. Little is known about differences in the time-course of retrieval at these two sites – and previous work has often used electroencephalography (EEG), which may lack the spatial resolution to separately resolve signals from ATL and pMTG. MEG studies of ATL show early responses (from 120ms) that appear to reflect interactions between semantic representations and inputs (Clarke et al., 2011; Mollo et al., 2017), plus later responses (250-450ms) that are influenced by patterns of coherent conceptual retrieval across both modalities (Marinkovic et al., 2003) and multiple items (Halgren et al., 2002; Lau et al., 2013; Bemis & Pylkkänen, 2011). Moreover, a recent chronometric TMS study by Jackson et al. (2015) found that the critical time point of involvement for ATL was around 400ms (although this study did not manipulate the strength of association and thus cannot identify when semantic processing in ATL is critical for the efficient retrieval of more coherent concepts). An N400 response has also been localised to pMTG (Helenius, Salmelin, Service & Connolly, 1998; Halgren et al., 2002; Lau, Phillips & Poeppel, 2008). This N400 effect is greater for unexpected meanings (Brown & Hagoort, 1993; Maess et al., 2006), although it also responds to a wide variety of semantic and lexical manipulations (Halgren et al., 2002; Lau, Phillips & Poeppel, 2008). In line with the N400 literature, ATL and pMTG can show a similar response to violations of semantic expectations – i.e., a stronger response to unrelated than related items (for a review, see Lau et al., 2008) – and thus the N400 semantic priming effect does not readily distinguish between ATL and pMTG; however, research has linked ATL to relatively automatic semantic priming (Lau et al., 2013) and the response in pMTG to more controlled or strategic semantic priming (Gold et al., 2006). E/MEG work has also shown that the response to unexpected meanings corresponds to a decrease in oscillatory power in the beta band, suggesting that oscillatory activity in this frequency range might support the maintenance of an appropriate network for comprehension given current expectations (Luoa et al., 2010; Wang et al., 2012; Kielar et al., 2014; Lewis & Bastiaansen, 2015).

Here, we used two temporally-sensitive methods (MEG, chronometric TMS) to examine the engagement of ATL and pMTG in semantic retrieval through time. By manipulating the strength of association between two words during explicit semantic decisions, we were able to test predictions of the Controlled Semantic Cognition framework (Lambon Ralph et al., 2017; Jefferies, 2013). By this account, pMTG is expected to show a stronger response to weak compared with strong associations, consistent with a role in controlled aspects of semantic retrieval, while ATL is predicted to show a stronger response to items coherent with dominant aspects of knowledge (i.e., effects of strong > weak associations). We were also able to test two alternative hypotheses about the timing of these effects. By one view, information is first retrieved and then selected: this account might envisage effects of strong associations in ATL that precede the engagement of controlled retrieval in pMTG. Alternatively, controlled retrieval processes in pMTG may be engaged at an early stage to shape patterns of semantic retrieval in ATL. In Experiment 1, we used beamforming analyses to characterise changes in total oscillatory power in ATL and pMTG during the retrieval of strong and weak associations. Total power includes components that are not phase-locked to an event/stimulus (i.e., responses that are generated at a slightly different time point across trials or participants). These so-called “induced” responses might be prominent in the retrieval of semantic relationships that span successive items (since the emergence of relationships between inputs might not be time and phase-locked to the onset of the second word). In Experiment 2, chronometric TMS was used to determine the causal role that anterior and posterior regions of the temporal lobe play in the retrieval of strong and weak associations at different time points. Together these two experiments, using different neuroscientific techniques, characterise the spatiotemporal basis of semantic retrieval when information is coherent with strongly-encoded aspects of knowledge, and show how this changes when non-dominant aspects of knowledge are required.

## Experiment 1: MEG

### Materials and Methods

#### Participants

Participants were 20 right-handed native English speakers, with normal or corrected-to-normal vision, and no history of language disorders (14 female, mean age 23.3 years, range 20-35). Data from one participant was excluded because their accuracy in the task fell below the acceptable minimum of 75% correct. Written consent was obtained from all participants and the study was approved by the York Neuroimaging Centre Research Ethics Committee.

#### Materials

The task and stimuli were adapted from Badre et al. (2005). Word pairs were presented, one word at a time, with varying associative strength between the first and second word, and participants were asked to decide if the two words were related in meaning or not. Participants were presented with 440 word pairs that were strongly-related (n=110), weakly-related (n=110), or unrelated (n=220). Strong and weak word pairs were selected using free association response data from the Edinburgh Associative Thesaurus (EAT). Strong associates were produced relatively frequently by participants (23%), while weak associates were produced more rarely (1%). The difference in mean association strength between strong and weak conditions was highly significant (t(188)=16.05, *p*<.001; Table 1). While our analyses focussed on the second word in each pair (which were identical across conditions between subjects), Table 1 confirms that there were no significant differences in word frequency or length across strong and weak conditions for the initial word.

**Table 1:** Comparing word frequency and length for the first word across conditions, plus the associative strength between the two words in the MEG experiment

| Measure | Strong Association | Weak Association | p-value |
| --- | --- | --- | --- |
|  | M (SD) | M (SD) |  |
| Word frequency | 26.6 (64.20) | 29.1 (38.0) | .59 |
| Word length (letters) | 5.5 (1.80) | 5.0 (1.5) | .16 |
| Association strength | 0.23 (0.19) | 0.01 (0.005) | .001 |

Unrelated trials were created by randomly shuffling words across pairs and manually removing any associations arising by chance. Target words were presented *either* following a strong or weak associate (not both), and in the unrelated condition. This meant that there was a 50% chance on any trial that a pair of words was semantically related.

#### Procedure

An illustration of the procedure can be seen in Figure 1a. Nonius lines (acting as a fixation cross) were present at all times. Before each trial, there was a rest period of 800 ms, plus an unpredictable jittered interval from 0 to 1000 ms, designed to reduce anticipatory responses. The first word was presented for 200 ms, there was an inter-stimulus interval (ISI) of 150 ms, and then the second word appeared for 200 ms followed by a 1000 ms interval. After each trial, the nonius lines were dimmed (for 1200 ms) and participants were encouraged to confine blinking to this period. The task required participants to make an explicit judgement about the relationship between the two words. On 10% of the trials, participants were cued to make an overt response by the presence of a question mark (on screen for 1000ms). They pressed one of two buttons with their left hand to indicate whether they had identified an association. These ‘catch trials’ were used to monitor performance in the task, and were excluded from further analysis. Since we only collected behavioural data for a small number of trials during MEG (to keep participants attending to the task), we also ran a behavioural version of the experiment outside the scanner, with the same participants, a minimum of 4 weeks before MEG data collection. This experiment was identical to the MEG version, except a response was given on every trial, and the pairings between stimuli were reversed – if a particular target was paired with a strong associate in the behavioural experiment, it was presented following a weak associate in MEG (and vice versa). Data from the behavioural experiment and the catch-trials in MEG are shown in Figure 1b and c.

**Figure 1:**
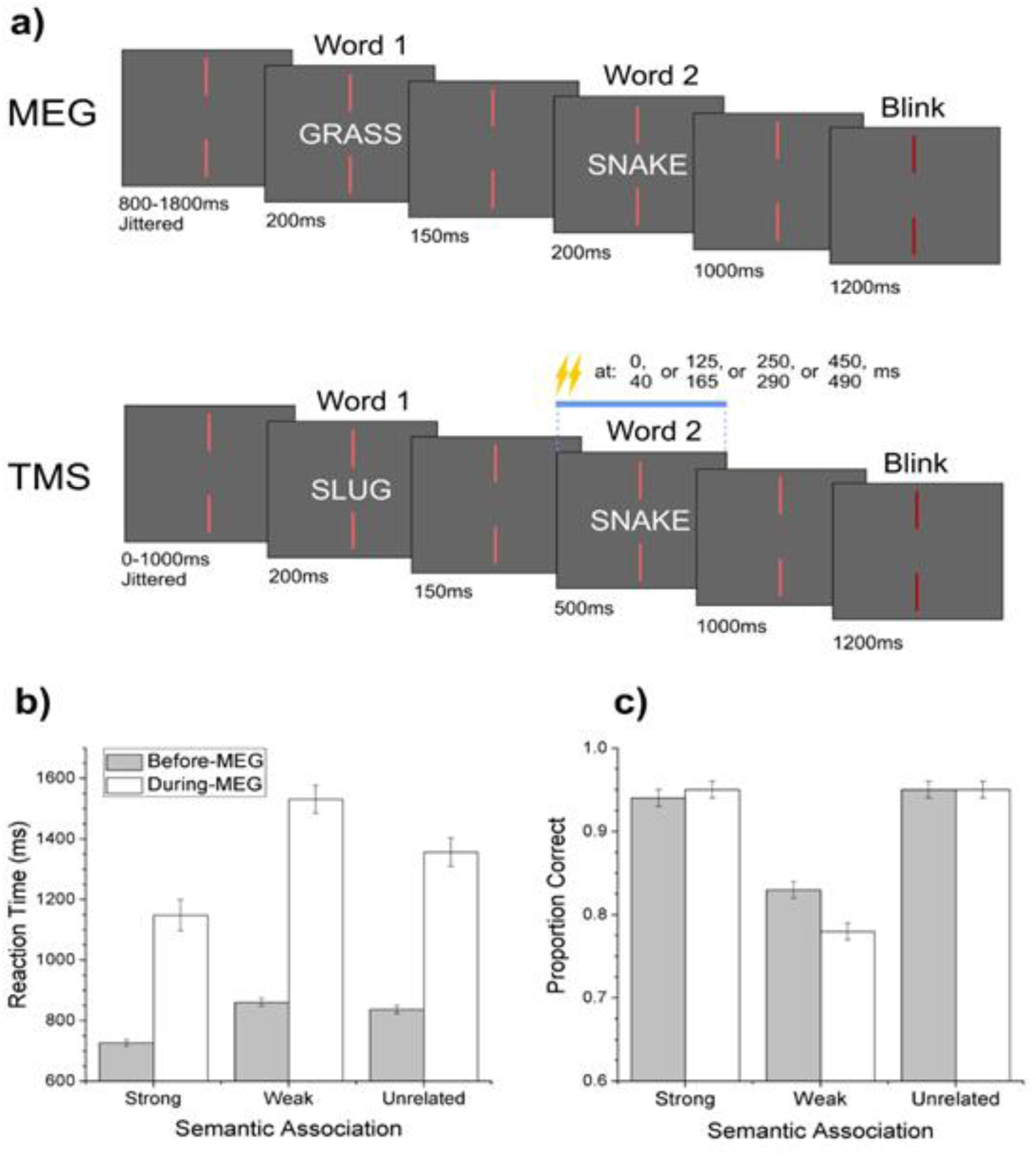
a) Example trials and timelines for the MEG and TMS experiments. b) Reaction time and c) accuracy data from the behavioural (gray bars) and MEG (white bars) experiments. Standard errors are corrected for repeated measures.

#### Stimulus presentation

The MEG experiment was carried out in a dark, magnetically shielded room. Presentation version 16.1 (Neurobehavioral Systems) was used to present the stimuli, communicate their timings to the MEG data acquisition system and to record participants’ responses on catch trials. Stimuli were back-projected onto a screen with a viewing distance of ~75 cm, so that letter strings subtended ~1° vertically and ~5° horizontally at the retina. Light grey letters on a dark grey background were used, such that the screen luminance was in the mesopic range, and a neutral density filter was used to minimize glare.

#### Data collection

During MEG recordings, participants were seated in an upright position, with the magnetometers arranged in a helmet shaped array, using a whole-head 248-channel, Magnes 3600 system (4D Neuroimaging, San Diego, California). Data were recorded in continuous mode, with a sampling rate of 678.17 Hz and pass-band filtered between 1-200 Hz.

Before MEG data acquisition, participants’ head shape and the location of five head coils were recorded with a 3D digitizer (Fastrak Polhemus). The signal from the head coils was used to localize the participant’s head position with respect to the magnetometer array before and after the experiment. The 3D digitized head shape of each participant was used for the co-registration of individual MEG data onto the participant’s structural MRI image using a surface-based alignment procedure from Kozinska, Carducci, and Nowinski (2001). For each participant, a high-resolution structural T1-weighted anatomical volume was acquired in a GE 3.0 T Signa Excite HDx system (General Electric, USA) at the York Neuroimaging Centre, University of York, with an 8-channel head coil and a sagittal-isotropic 3-D fast spoiled gradient-recalled sequence (repetition time/echo time/flip angle = 8.03 msec/3.07 msec/20°, spatial resolution of 1.13 mm × 1.13 mm × 1.0 mm, in-plane resolution of 256 × 256 × 176 contiguous slices).

External, non-biological noise detected by the MEG reference channels was subtracted, and MEG data were converted into epochs of 1500 ms length, starting 800 ms before the onset of the first word. All channels from all trials were inspected visually and epochs containing artifacts, such as blinks, articulatory movements, swallowing, and other movements, were rejected manually. Data from three malfunctioning channels were automatically rejected for all participants. Statistical analyses included only datasets with at least 75% of trials retained after artefact rejection. 20 (of 21) datasets reached this criterion. On average, 17% of the trials were rejected from these datasets (min 7.3% - max 25%).

#### MEG analysis strategy

The first task was to identify points of interest (POIs) in ATL and pMTG showing strong oscillatory responses to the experimental manipulations in general. To achieve this, we carried out a whole-brain analysis of all the related trials, collapsed across the strong and weak conditions, at coarse frequency and time resolution. Secondly, the fine-grained time-frequency responses of these POIs were compared across experimental conditions. This analysis strategy allowed us to assess the contributions of ATL and pMTG, indexed by their time-frequency responses, in relation to the Controlled Semantic Cognition hypothesis (Jefferies, 2013; Lambon Ralph et al., 2017). For both whole-brain and POI analyses, sources of neural activity were reconstructed with a modified version of the vectorised, linearly-constrained minimum-variance (LCMV) beamformer (Van Veen et al., 1997; Huang et al., 2004), implemented as part of the public domain Neuroimaging Analysis Framework (NAF) pipeline at the York Neuroimaging Centre (http://vcs.ynic.york.ac.uk/docs/naf/index.html). We used a multiple spheres head model (Huang et al., 1999), with co-registration checked manually. An MEG beamformer (spatial filter) allows the signal coming from a location of interest in the brain to be estimated while attenuating signals from elsewhere. This is achieved by reconstructing the neuronal signal at a specific point (referred to as a Virtual Electrode) as the weighted sum of the signals recorded by the MEG sensors. The covariance matrix, used to generate the weights for each beamformer, was regularized using an estimate of noise covariance (Prendergast et al., 2011; Hymers et al., 2010). This procedure was performed separately for each condition and/or analysis window, in order to optimise sensitivity to the effect of interest (Brookes et al., 2008; 2011). The outputs of the three spatial filters at each point in the brain were summed to generate estimates of oscillatory power. This analysis strategy and the parameters used for the current study were similar to those used in recent MEG studies of visual word recognition, object naming and semantic processing (Wheat et al., 2010; Klein et al., 2014; Urooj, 2014; Mollo et al., 2017).

##### Whole brain beamforming

The brain’s overall response to the task (collapsing the strong and weak trials) was characterised within broad frequency ranges and relatively long periods of 200ms. A cubic lattice of points was defined in the brain (5 mm spacing), and at each point, an independent set of spatial filters was defined to estimate the source current at that point. A noise-normalised volumetric map of total oscillatory power (i.e., including both the evoked and non-phase locked components) was then produced over these broad temporal windows and frequency bands. Since our main research question concerned how the brain’s response to the second word changed as a function of its relationship to the first word, we defined time zero as the onset of the second word of the pair; the onset of the first word was at −350ms relative to this. We then characterised whole-brain oscillatory responses to the second word by contrasting responses in “active” time windows at 0-200ms, 200-400ms, and 400-600ms post-onset of the second word with a 200ms “passive” time window at −700 to −500ms (prior to the onset of the trial). The Neural Activity Index (NAI; Van Veen et al., 1997), which is an estimate of oscillatory power that takes account of spatially-inhomogeneous noise, was calculated at each point in the lattice, within the following frequency pass-bands: 5-15 Hz, 15-25 Hz, 25-35 Hz and 35-50 Hz. These frequency ranges were taken from previous MEG studies of reading (Klein et al., 2014; Wheat et al., 2014). This analysis produced an NAI volumetric map for the active and passive period, separately for each participant at each frequency band, from which paired-samples t-statistics were calculated. Individual participant’s t-maps were then transformed into the MNI standardized space in order for group level statistics to be calculated. To do this, a null distribution was built up by randomly relabelling the active and passive windows for each participant at each grid point, using the permutation procedure developed by Holmes et al. (1996). The maximum t-value obtained with random relabelling across 10000 permutations was established. We then compared the real distribution of t-values in the data with the maximum t-value obtained from the permuted data. Maximum statistics can be used to overcome the issue of multiple comparisons (i.e. controlling experiment-wise type I error), since the approach uses the highest permuted t-value across the brain to provide a statistical threshold for the whole lattice of points, over which the null hypothesis can be rejected (Holmes et al., 1996). Figure 2 shows those areas in the brain with t-values equal or higher than the top 5% of t-values present in the null distribution.

**Figure 2:**
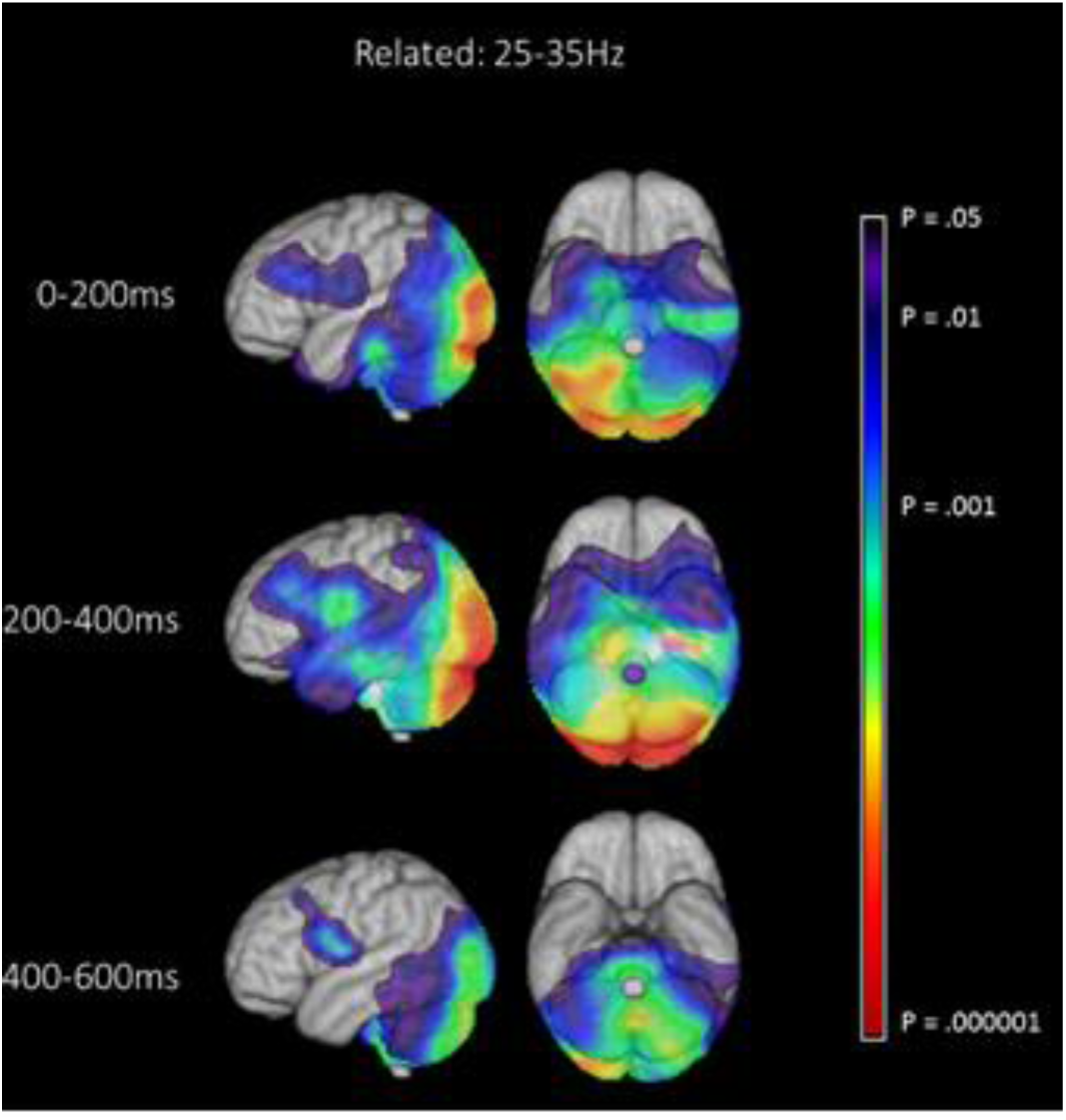
Whole-brain beamforming results for the 25-35 Hz frequency band, showing differences in total oscillatory power between an active period following target onset and a passive period prior to each trial. The first 600ms following presentation of target word are displayed, in 200ms windows. Task effects were decreases in total power in all cases. The images show a t-value map, thresholded at p<.05.

##### Time-Frequency Analysis: Point of Interest (POI)

In the whole brain analysis, oscillatory signals were strongest and most extensive in the 25-35Hz frequency band, within the 200-400ms time window, particularly in ATL and pMTG (see Figure 2) and therefore peaks in these maps were used to identify POIs. Other frequency bands showed a similar but not as extensive a distribution of oscillatory power (see Supplementary Materials Figure 1). Following the selection of POIs from the whole-brain beamforming analysis, separate beamformers (Huang et al., 2004) were used to reconstruct the neural activity in ATL (MNI coordinates −48,8,−18) and pMTG (MNI coordinates −50,−52,8). These sites corresponded to points showing task-induced changes in oscillatory power within the relevant regions of cortex. Although bilateral ATL is implicated in semantic representation, we focused on left-hemisphere sites since (i) the stimuli were written words; (ii) fMRI and patient studies reveal a greater contribution of the left hemisphere to semantic processing, especially for written words (Binder et al., 2009; Rice, Lambon Ralph & Hoffman, 2015; Noonan et al., 2013); and (iii) right motor cortex was expected to show irrelevant responses related to the preparation of button presses with the left hand (even though button presses were only required on catch trials), and therefore contaminate the signals of interest.

We then used the Stockwell transform (Stockwell, Mansinha, & Lowe, 1996) to calculate time-frequency representations for each POI from 5-50 Hz over the time period −800 to 700 ms, where 0ms was the onset of the second word. This allowed us to examine the response to semantic matching from 0-600ms, with reference to a passive period before the onset of the first word (defined as −700 to −500 ms as in the whole brain analysis). The Stockwell transform, implemented in the NAF software, uses a variable window length for the analysis which is automatically adapted along the frequency range according to the sample rate and the trial length (4^th^ order Butterworth filters with automatic padding). The time-frequency representations of total power were normalized, separately for each condition and for each participant, with respect to the mean power per frequency bin in a baseline period prior to the start of trials in that condition (−700 to −500 ms). This window length was also used in earlier studies (Mollo et al., 2017; Wheat et al., 2010; Klein et al., 2014), since it provides a compromise between the minimum length sufficient to estimate power at the lowest frequency reported here (i.e., 5Hz) and the requirement to characterise the state of the brain immediately before the onset of each trial.

To compare the time-frequency representations between experimental conditions, we used PROC MIXED in SAS (SAS Institute Inc., North Carolina, US) to compute generalized linear mixed models (GLMM). Time-frequency plots of percentage signal change were treated as two dimensional arrays of small time-frequency tiles, indexed in the model by three main effects: time, frequency and the interaction between time and frequency. Therefore, random effects were included in each GLMM to account for the fact that each participant’s time-frequency plot is made up of multiple time-frequency tiles. Time-frequency (or spatial) co-variance in the spectrogram was controlled for by assuming the estimates of power followed a Gaussian distribution: consequently a Gaussian link function was used in the model. The time-frequency (spatial) variability was integrated in the model by specifying an exponential spatial correlation model for the model residuals (Littel et al., 2006). Finally, the data were resampled at a frequency resolution of 2.5Hz and time resolution of 25ms, the smallest time and frequency bin consistent with model convergence. This time-frequency resolution proved optimal in other similar published studies (Mollo et al., 2017; Klein et al., 2014; Urooj et al., 2014; Wheat et al., 2010). PROC MIXED constructs an approximate *t* test to examine the null hypothesis that the LS-Mean for percentage signal change between conditions was equal zero in each time-frequency tile, and the procedure automatically controls for multiple comparisons (i.e. controlling experiment-wise type I error). The statistical contours on the percentage signal change figures for total power encompass time-frequency tiles fulfilling both of the following criteria: a) the difference between conditions reached *p* < 0.05; b) any region in the time-frequency plot defined by (a) also showed a response that was significantly different from zero in at least one of the two contributing conditions.

## Results

### Behavioural experiment

While traditional priming experiments show facilitation for weakly-related as well as strongly-related primes, compared with unrelated words (Neely, 1977; 1991), weak associations are expected to show a processing cost when making explicit semantic decisions, since semantic control is expected to be required to promote information relevant to the specific association being probed and to suppress dominant but currently-irrelevant features (Badre et al., 2005; Whitney et al., 2011). The behavioural data were consistent with these predictions (Figure 1b and 1c). A one-way repeated-measures ANOVA of reaction times from the behavioural pre-scan results showed a statistically significant main effect of experimental condition (F(2,38) = 22.26, p<.001; Figure 1b). Post-hoc comparisons showed that reaction times were faster for strong associations compared with both the weak and unrelated conditions (t(38)=6.25, p<.001 and t(38)=5.15, p<.001 respectively). There was no statistically significant difference in reaction times between the weak and unrelated conditions. A similar analysis for accuracy showed a statistically significant main effect of condition (F(2,38) = 31.47, p<.001; Figure 1c). Post-hoc comparisons showed that accuracy for the weak condition was significantly lower than that for both the strong and unrelated conditions (t(38)=6.78, p<.001 and t(38)=6.95, p<.001). There was no significant difference in accuracy between the strong and unrelated conditions within pre-scan behavioural experiment.

Reaction times were generally longer for catch-trials recorded during MEG acquisition yet the data followed a similar pattern to the pre-scan experiment. A one-way repeated-measures ANOVA of reaction times from the catch-trials in MEG showed a statistically significant main effect of experimental condition (F(2,38)=10.63, p<.001), as shown in Figure 1b. Post-hoc comparisons showed faster reaction times for strong associations compared with both the weak and unrelated conditions (t(38)=4.60, p<.001 and t(38)=2.50, p<.05 respectively). In addition, reaction times for the unrelated condition were significantly faster than those for the weak condition (t(38)=2.10, p<.05). A similar analysis of catch-trial accuracy showed a main effect of condition (F(2,38)=89.03, p<.001), as shown in Figure 1c. Post-hoc comparisons showed that accuracy for the weak condition was significantly lower than that for both the strong and unrelated conditions (t(38)=11.66, p<.001 and t(38)=11.45, p<.001). However, there was no significant difference in accuracy between the strong and unrelated conditions.

### Whole-brain results

The response to the task as a whole (i.e., the response to the second word of the pair collapsed across both experimental conditions, versus a period prior to the start of the trial), is shown in Figure 2. The most extensive changes in total power in response to the task were power decreases, relative to the resting passive period, in the 25-35Hz frequency band (shown in Figure 2 below), although other frequency bands showed similar effects (see supplementary Figure 1). These decreases in total oscillatory power were focussed on temporal, occipital, inferior frontal and parietal lobe regions implicated in visual and semantic processing, starting within the first 200ms and lasting for at least 600 ms after target presentation. Decreases in total power are commonly interpreted as reflecting an *increase* in neural activity that is not phase-locked to stimulus presentation (Hanslmayr et al., 2012). Reductions in total power have been shown to correlate with an increased BOLD response in fMRI (Hanslmayr et al., 2011; Singh et al., 2002; Hall et al., 2014), and a recent review proposed that decreases in total power reflect active engagement of neocortex in the encoding and retrieval of memories (Hanslmayr, Staresina & Bowman, 2016). Thus, the whole-brain beamforming results are consistent with an increase in visual and semantic processing following the onset of the second word.

### Points of interest results

#### Whole epoch data for each site

For each POI, Figure 3 shows time-frequency plots of total power for the whole epoch, corresponding to the first and second word responses in each semantically-related pair. These plots are included to illustrate the response to the task at each site, and to inform the interpretation of contrasts between conditions that were computed from the onset of the second word, in the context of ongoing task activity. Orange-red-brown colours indicate *power increases*, whereas green-purple-black colours indicate *power decreases* relative to the baseline (with no change shown in green). In both ATL and pMTG, there was a subtle increase in oscillatory power in response to the first word, while the presentation of the second word was characterised by a large *reduction* in total oscillatory power relative to baseline. This effect followed the offset of the first word (and was particularly pronounced for pMTG); the decreases in total power then became stronger and encompassed more frequency bands in response to the onset of the second word.

**Figure 3:**
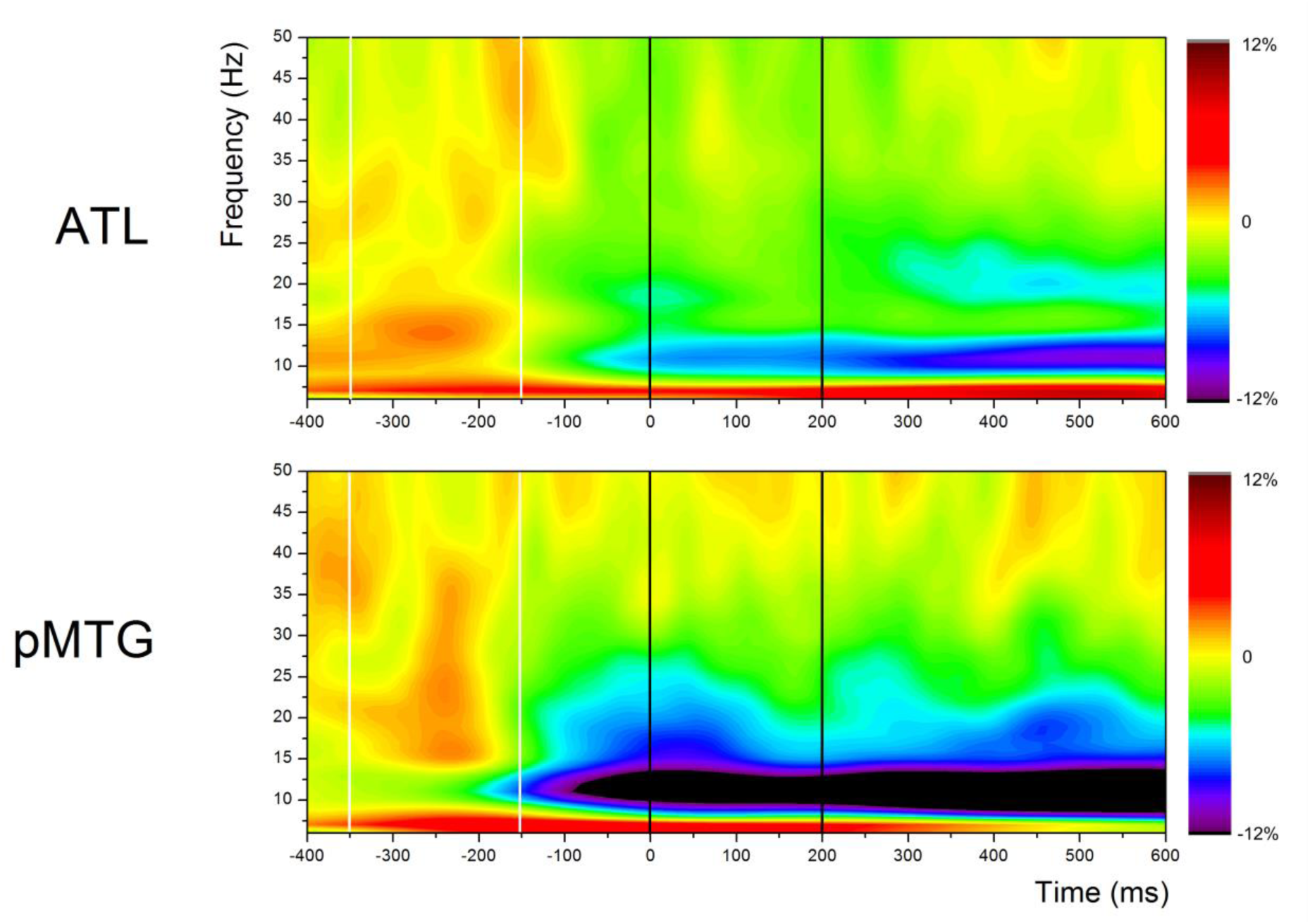
Total oscillatory power across the whole epoch for related trials, including both words presented in the relatedness judgement task. Presentation of the prime word (first word of the pair) is shown within white vertical lines, while presentation of the target word (second word of the pair) is illustrated within black vertical lines. Orange-brown indicates regions of *power increase* relative to the baseline, while green-purple indicates *power decreases* relative to the baseline, and yellow indicates no change from baseline

#### Differences between conditions in POIs

As shown in Figure 4c, we found statistically significant differences between strong and weak associations throughout the epoch, in the beta and low gamma frequency bands, in both ATL and pMTG. However, strength of association had opposite effects at the two sites, in line with the predictions of the Controlled Semantic Cognition framework (Lambon Ralph et al., 2017). ATL showed a greater change from baseline during the retrieval of strong vs. weak associations. This effect was significant from 400ms post-target onset until the end of the epoch at 7-12 Hz. PMTG, in contrast, showed stronger changes in oscillatory power during the retrieval of weak associations, from within 100ms of the onset of the second word, and this effect lasted throughout the epoch (to 550ms, at around 15Hz, plus brief significant differences at 25Hz and 30Hz).

**Figure 4:**
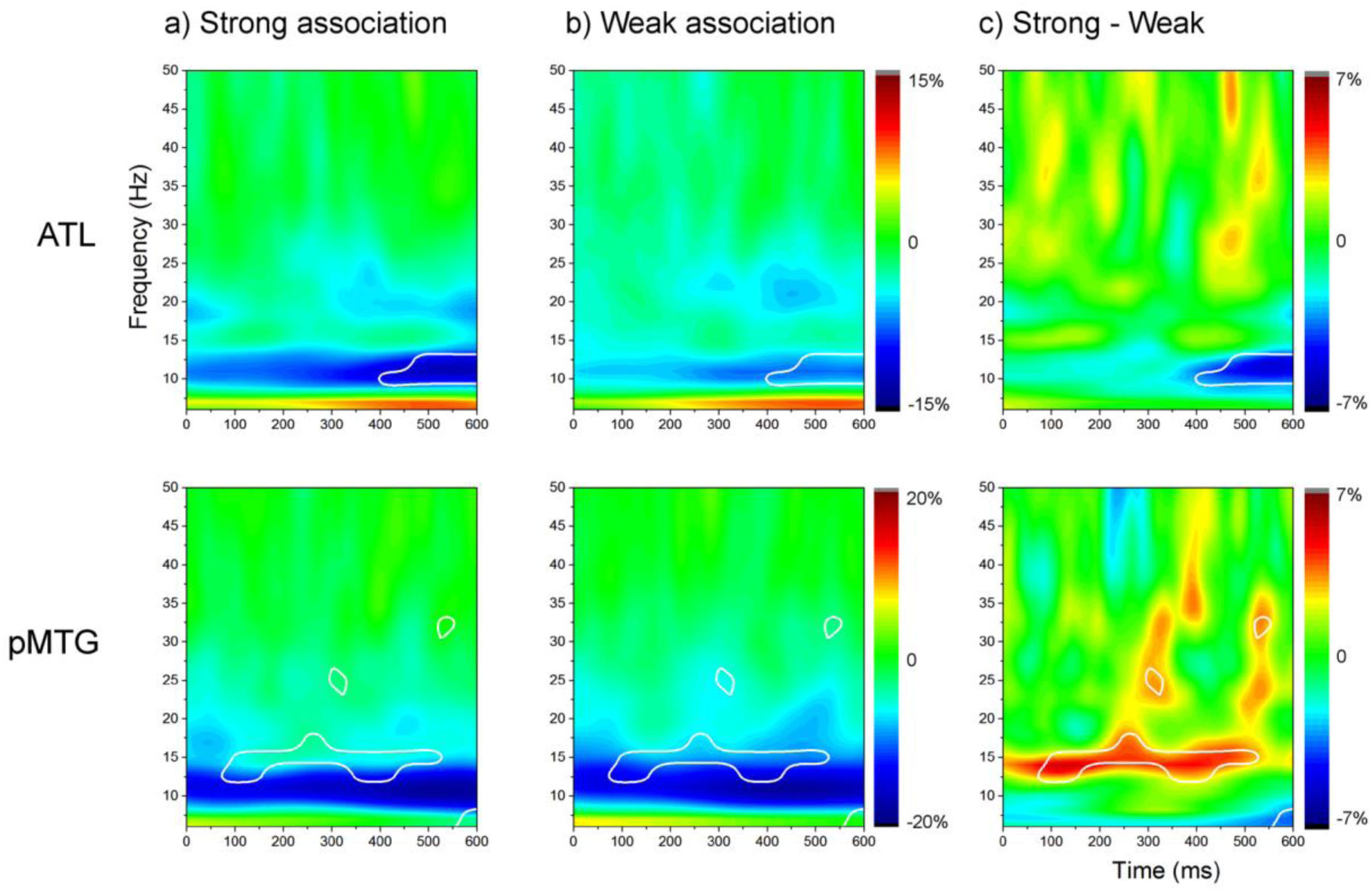
a) Percentage signal change in the strong condition, relative to baseline. b) Percentage signal change in the weak condition, relative to baseline. c) Percentage signal change between strong and weak conditions, separately for ATL and pMTG. White lines are derived from the statistical comparison between strong and weak conditions. The boundaries enclose regions fulfilling two criteria: i) percentage signal change between the strong and weak conditions is significantly different from zero (p<0.05) and ii) percentage signal change computed separately for each condition is significantly different from zero for at least one of the two conditions. Yellow-red colours indicate regions of *power increase* relative to the baseline, while cyan-blue indicates *power decreases* relative to the baseline, and green indicates no change from baseline.

While the focus of this study was on differences between strong and weak associations to test the predictions of the Controlled Semantic Cognition Framework (Lambon Ralph et al., 2017), we also computed differences between related and unrelated trials to allow comparison with previous studies that employed similar contrasts (for a review, see Lau et al., 2008). The results of this analysis can be seen in Supplementary Materials.

#### Summary of MEG results

Comparisons of strong and weak associations revealed a dissociation in the temporal lobe, in both space and time, which depended on the match between the semantic retrieval required by the task and the structure of long-term conceptual knowledge. ATL showed a strong response during the retrieval of both strong and weak associations soon after the presentation of the second word, plus greater oscillatory power for strong than weak associations from around 400ms after target onset. This is consistent with the view that ATL supports coherent semantic retrieval when inputs and task requirements align with long-term conceptual representations (Feng et al., 2016; Binder, 2016). The timing of this result is consistent with previous studies showing strong semantic effects in ATL around 400ms and suggests that effects of coherent semantic retrieval emerge over time (Jackson, Hoffman, Pobric, & Lambon Ralph, 2016; Lau et al., 2013; 2014; Marincovic et al., 2003). In contrast, pMTG showed greater oscillatory power for weak than strong associations soon after target onset and throughout the epoch, in line with its purported role in more controlled patterns of semantic retrieval. Since pMTG responded rapidly after the onset of the second word, this site might play a role in maintaining a semantic context from the first word and detecting circumstances where inputs are inconsistent with this context (triggering the recruitment of controlled retrieval processes). Given the sustained engagement of pMTG for weak > strong associations, this site might also play a role in shaping ongoing semantic retrieval such that it focuses on the aspects of knowledge that are currently relevant.

## Experiment 2: Chronometric TMS

Experiment 1 demonstrated a dissociation in oscillatory power within the temporal lobe in space and time, reflecting the extent to which the pattern of semantic retrieval required by the task was consistent with dominant aspects of long-term conceptual knowledge. To determine the causal role of ATL and pMTG in semantic retrieval, Experiment 2 used chronometric TMS to disrupt processing in these two regions at different points in time in the same paradigm. Stimulation of ATL during the presentation of the second word in the pair might disrupt the efficient retrieval of strong associations, given the MEG findings above, while stimulation of pMTG at the moment of onset of the second word might disrupt the retrieval of weak associations if this site supports the *engagement* of controlled retrieval processes.

### Materials and Methods

#### Participants

Participants were 15 right-handed native English speakers, with normal or corrected-to-normal vision, and no history of language disorders (8 males, mean age 23, age range 20-32 years). Written consent was obtained from all participants and the study was approved by the York Neuroimaging Centre Research Ethics Committee.

#### Design

The experiment employed a 3x2x4 repeated-measures design, with site (ATL, pMTG and sham mid-MTG), task (semantic association task and digit parity judgement task), and TMS timings (0 & 40ms; 125 & 165ms; 250 & 290ms; 450 & 490ms) as within-subject factors. At each time point, a pair of pulses 40ms apart were applied, since this dual-pulse method is thought to generate more significant behavioural disruption than single pulses (Gagnon, Schneider, Grondin & Blanchet, 2011; Strafella & Paus, 2001; Chen, 2000). The stimulation times were selected to provide coverage of time points of interest from the MEG experiment: these included processes in already play by the onset of the second word, which are likely to be important given the successive stimulus presentation used in our paradigm, responses observed 100-200ms after the onset of the second word (by which point the differential response in pMTG was established), effects within the first 300ms (e.g., related > unrelated differences in ATL), and later effects. This allowed us to explore these sites’ causal involvement in retrieving dominant and weaker aspects of knowledge.

#### Materials

The semantic task was the same as for Experiment 1. Word pairs were presented sequentially, and participants decided whether the two words were related or not. The pairs were either strongly or weakly associated, or they were unrelated. To maximise sensitivity to the effects of TMS on the retrieval of strong and weak associations, each session comprised 70% related trials (which were the focus of the analysis) and 30% unrelated trials (to ensure participants attended to the task, which were excluded from the analysis). The same target words were presented across conditions, although each target was only presented once per session. In addition, the first words of the strong and weak pairs did not differ in word frequency or length (see Table 2).

**Table 2:** Comparing word frequency and length for the first word across conditions, plus the associative strength between the two words in the TMS experiment

| Measure | Strong Association | Weak Association | p-value |
| --- | --- | --- | --- |
|  | M (SD) | M (SD) |  |
| Word frequency | 17.43 (32.38) | 19.28 (32.91) | .66 |
| Word length (letters) | 5.62 (1.81) | 5.48 (1.51) | .49 |
| Association strength | 0.43 (0.19) | 0.03 (0.06) | .001 |

A non-semantic task involving numerical judgements was designed to match the semantic task in overall difficulty. Two three-digit numbers were presented sequentially, and subjects were asked to decide whether *both numbers* were odd or even. The proportion of yes/no trials was identical to the semantic task (i.e., 70% match). One participant was tested on a different number judgement task and was excluded from the statistical comparisons of semantic vs. number task performance. For the word conditions, there were 25 trials with TMS delivered to each of the three stimulation sites at 4 different timings (25×4×3), for each condition (strongly related, weakly related, unrelated). For the digit task, there were 12 trials for each of the three stimulation sites at 4 different timings (12×4×3), for each number “condition” (both even, both odd, different).

#### Stimulus presentation

The three experimental sessions were divided into 5 runs, each lasting approximately 12 minutes. TMS was delivered in 4 of the 5 runs, and a block without TMS was placed in the middle of the 5 runs for safety reasons. Each run was made up of 6 blocks for each task (numerical or semantic), lasting around 60 seconds. Blocks were arranged in pseudorandom order to minimise task switch costs. When switching between tasks, a short instruction screen informed the participant which task would be presented next. The first trial after the task switch was a dummy trial which was discarded from further analysis. The first word of the pair was presented for 200ms, followed by an inter-stimulus interval (ISI) of 150ms, and then the second word requiring a relatedness judgement appeared for 500ms (see Figure 1a). The nonius lines remained on screen for 1000ms, and were then dimmed for 1150ms after the participant’s response, to signal the end of the trial. Following this, the bright nonius lines returned, to cue the onset of the next trial, for a randomly variable interval of 0-1000ms (500ms on average) before the onset of the first word of the next pair. Each trial lasted on average 3500ms. As in the MEG experiment, participants were asked to decide if the two words were related in meaning or not. They responded with their right hand and were instructed to be as quick and accurate as possible. Before starting the experiment, participants performed a practice session with 10 trials of both tasks (without TMS), and three practice trials with stimulation. Participants took self-paced breaks between the runs.

#### Stimulation sites

TMS was applied to left ATL, left pMTG, and a sham site in the mid-temporal lobe (halfway between these two sites). Stimulation sites were taken from published studies; participants’ structural T1 MRI scans were co-registered to the scalp using the Brainsight frameless stereotaxy system (Rogue Research, Montreal, Canada) to identify the stimulation targets in each participant’s brain. The left ATL site was in anterior ventrolateral temporal cortex (MNI −51, 6, −39; coordinates from Binney et al., 2010). This site showed greater activation for synonym judgement than numerical magnitude judgement in fMRI, and is located close to the region of peak atrophy in semantic dementia (Binney et al., 2010). The left ATL coordinates for TMS fell within the area of statistically significant oscillatory power revealed by whole-brain beamforming in Experiment 1, although the peak in the MEG data was anatomically superior (21 mm), and somewhat lateral and anterior (3 mm and 2 mm respectively) relative to the stimulation site. Similarly, the choice of left pMTG site for TMS was based on a metaanalysis of neuroimaging studies of semantic control by Noonan et al. (2013; MNI −58, −50, −6). This site activates across a wide range of manipulations of semantic control, and shows a stronger response to weak than strong associations (Davey et al., 2016; Gold et al., 2006). It was also located within the area of statistically significant oscillatory power revealed by whole-brain beamforming in Experiment 1, but was inferior (14 mm) and lateral (8 mm) to the pMTG POI. We opted to use stimulation sites from the literature rather than peaks from Experiment 1 given the relatively poor spatial resolution of MEG. The sham control site was selected by finding the midpoint on the y-axis between the two experimental sites, varying the z coordinate to ensure that stimulation was delivered to the middle temporal gyrus.

#### TMS stimulation protocol

Chronometric TMS was delivered using a Magstim Rapid2 stimulator and a 50mm diameter figure-eight coil. Stimulation intensity for ATL and pMTG was 60% of the maximum output of the stimulator. Sham stimulation was applied at 30% of stimulator output since this intensity is thought to be too weak to produce a neural effect, but it still mimics the sound and scalp sensations of TMS stimulation (Duecker et al., 2013). Dual-pulse TMS was delivered at 25Hz (pulses 40 ms apart) in each trial (see Figure 1a for illustration). The position of the coil was monitored and tracked in real time. The mean difference between the intended target and the stimulated site on each trial was 0.3mm (s.d. = 0.26; maximum displacement = 5.6mm). Trials in the different timing conditions were arranged in an ascending or descending staircase of 4 trials (i.e. four trials with stimulation at 0 & 40ms followed by four trials of stimulation at 125 & 165ms etc.). We used this strategy to limit the participants’ awareness of the different TMS timings, and to reduce any tendency to wait until stimulation had been delivered before responding (Sliwinska et al., 2012). Following safety guidelines (Rossi et al., 2009), an inter-train interval of 5000ms was added after every sequence of 24 double pulses. Where possible this interval corresponded to the task switching instruction screen; in other cases it was added after a button press response.

#### Analysis strategy

We wanted to know how speeded judgements about strong and weak semantic relationships between pairs of words would be affected by TMS, delivered at different time points following the onset of the second word in a pair, at the two different cortical sites. To maximize the sensitivity of these analyses, we used generalised linear mixed models (GLMM) which retained information about all trials and permitted random effects at both the participant and item levels to be modelled (see Baayen, Davidson & Bates, 2008). To do this, we specified an ‘unstructured’ variance-covariance structure for each random effect in the model’s G-matrix. The mixed models were implemented in PROC MIXED in SAS v9.4 (SAS Institute, North Carolina, USA).

Previous TMS studies have reported consistent slowing for semantic decisions following inhibitory stimulation, and little effect on accuracy (Walsh & Cowey, 2000; Pasqual-Leone, Walsh & Rothwell, 2000; Devlin, Matthews & Rushworth, 2003). Therefore, our primary outcome variable for each trial was the magnitude of the TMS effect, defined as the difference in response time between a word pair subject to TMS and its corresponding sham version. Incorrect responses and outlying data points that fell more than 2SD from each participant’s mean RT were removed, for each session, prior to analysis.

For the initial models, we included the main effects of task condition (e.g., strong vs. weak association), site (ATL, pMTG), and TMS time (i.e., pulses at 0-40ms; 125-165ms; 250-290ms; 450-490ms after the onset of the second word), plus their interactions. We also included as covariates structural aspects of the experiment (i.e. session and block order). In addition, supplementary analyses, characterising (i) the effect of TMS on accuracy for strong and weakly-related targets and (ii) the effect of TMS on semantic judgements overall (vs. numerical judgements), highlighted non-specific effects of TMS on both RT and accuracy in our data, as we report in the Supplementary Materials. For these reasons, accuracy per block and performance in the numerical task were also included as covariates in the initial models for reaction time. The criteria we used to optimize the final model were: (i) a significant reduction in −2Log-Likelihood relative to the empty model, (ii) only explanatory variables that were statistically significant at p<.05 should be retained. Once the final model was fitted, we used PROC MIXED to estimate pairwise t-test comparisons of the least squared (LS) mean reaction times, with and without TMS, carried out separately at each site for each condition (a total of 5041 observations). These post-hoc comparisons were controlled for multiple comparisons.

## Results

The main effects from the optimized GLMM of reaction time are shown in Table 3. Since our dependent measure was the TMS effect (computed as the difference between TMS and sham trials), there was no main effect of condition. We found significant main effects of TMS time (reflecting greater differences between stimulation and sham at particular time points) and site (reflecting a greater difference between ATL and sham than between pMTG and sham). The covariates of block, session order, and number RT (i.e., non-specific effects of TMS; see Supplementary Materials) were also statistically significant, although the accuracy covariate did not improve model fit and was not included in the final model. Critically, there was a significant three-way interaction between condition (strong vs. weak), TMS time and site, suggesting that the disruption of strong and weak associations occurred at a different point in time after the onset of the second word of the pair, and that this effect was different comparing ATL with pMTG.

**Table 3:** Effect of TMS on RT for strong and weak associations

| Model Parameter | F-value (d.f.) | Z-value | p-value | -2Log<br>likelihood |
| --- | --- | --- | --- | --- |
| Empty model |  |  |  | 63784.3 |
| Time | 4.86 (3, 302) |  | .0026 | 63277.5 |
| Site | 12.39 (1, 4783) |  | <.001 |  |
| Condition × Time × Site | 1.95 (11, 4920) |  | .029 |  |
| Block order | 15.16 (3,5004) |  | <.001 |  |
| Testing session | 4.41 (2, 4465) |  | .012 |  |
| Number task RT | 191.9 (1,1955) |  | <.001 |  |
| Participant covariance |  | 2.59 | .0048 |  |
| Target covariance |  | 4.75 | <.001 |  |

Figure 5 shows mean reaction times (upper row) and the post-hoc comparisons of LSmean reaction times (low row), separately for ATL and pMTG. For ATL, we found a significantly larger effect of TMS on strongly-related than weakly-related pairs (giving rise to a positive LSmean difference in the bottom row of Figure 5), when pulses were applied at 125-165ms after the onset of the second word. At the other time points, the magnitude of the TMS effect was equivalent for the strong and weak associations. This suggests that at around 150ms post-presentation of the second word, the efficient retrieval of strong semantic relationships was disrupted by the perturbation of ongoing processing within ATL. Although strong associations did not evoke a stronger change in oscillatory response at this site until later (400ms in the MEG data), and the behavioural response was later still (between 500-600ms in this experiment), disruption of a settling process within ATL might potentially disrupt or delay both of these subsequent effects.

**Figure 5:**
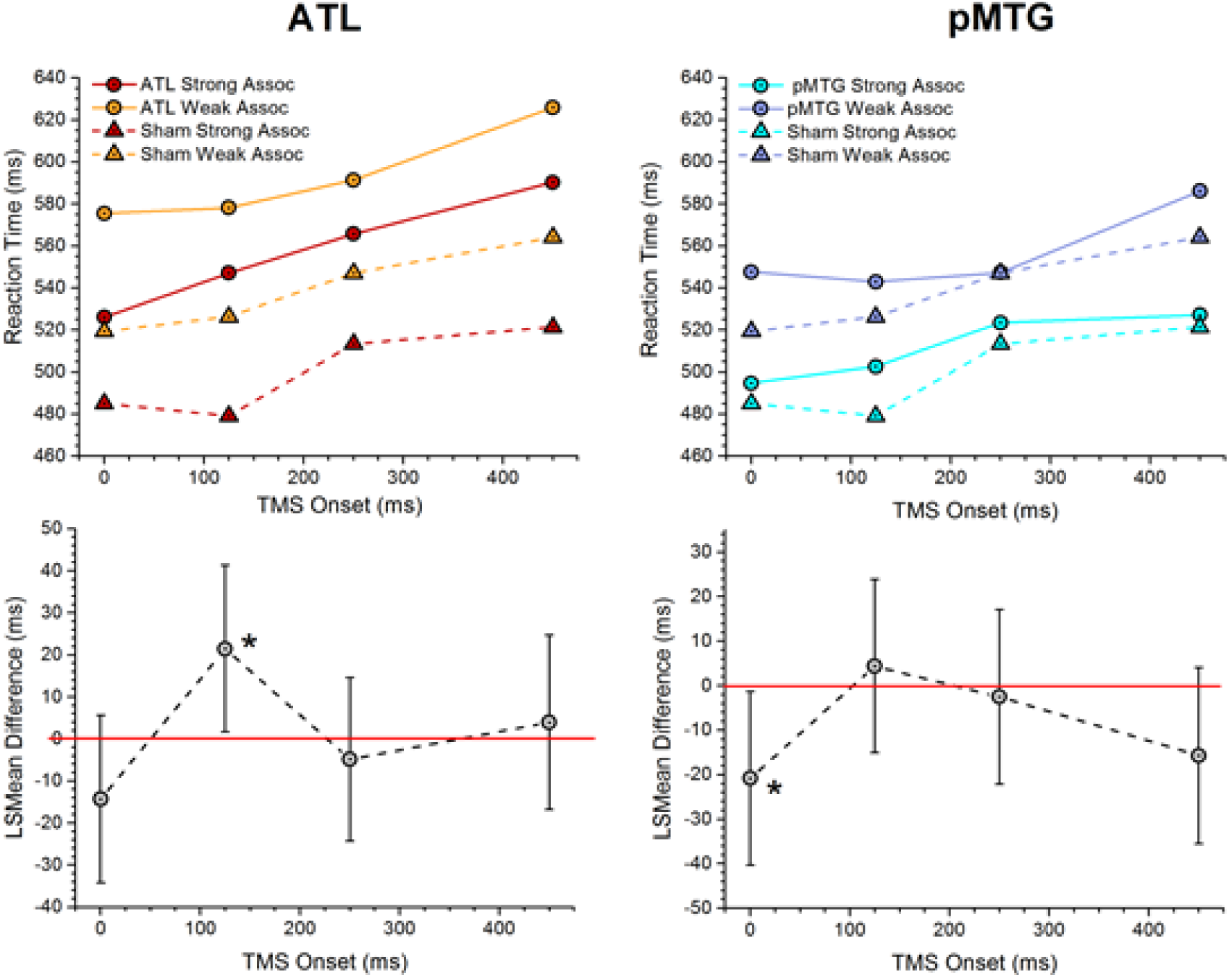
Effect of TMS on RT for strong and weak associations. TOP ROW: RT (in ms) for the strong and weak conditions for ATL (left) and pMTG (right). RT data for the strong and weak condition for the sham site is showed in dashed lines. These plots show the raw (un-modelled) means. BOTTOM ROW: A comparison of LS Means differences between strong and weak conditions in the effect of TMS. Data points above the red line indicate greater disruption for the strong condition, while data points below the red line indicate greater disruption for the weak condition. Statistically significant differences (at p<.05) between the effects of TMS on strong and weak trials are indicated with asterisks.

For pMTG, we found a significantly larger effect of TMS on weakly-related than strongly-related pairs (giving rise to a negative LSmean difference in the bottom row of Figure 5), when pulses were applied at 0-40ms after the onset of the second word in the pair. At the other time points, the magnitude of the TMS effect was equivalent for the strong and weak associations. This very early differential response suggests that pMTG may make a critical contribution to the capacity to *engage* controlled retrieval when it is needed. Stimulation at this early point may have disrupted the maintenance of current contextual information generated by the first word in the pair. This could disproportionately affect weak associations if, for example, pMTG plays a critical role in detecting the need to employ controlled retrieval. Although weak associations did not evoke a stronger change in oscillatory response at pMTG until slightly later (from around 60ms in the MEG data), effects linked to controlled retrieval at pMTG in both MEG and TMS were observed very early after the onset of the second word, allowing us to reject one view of the emergence of semantic retrieval over time, in which conceptual knowledge is first activated or retrieved and then subsequently selected to suit current task demands or the context.

## Discussion

A significant body of research has characterised the brain regions that support semantic processing but less is known about the temporal evolution of semantic retrieval across these regions. While studies have examined the time course of semantic access from written words and pictures following a semantically-related or an unrelated item (Dikker & Pylkkänen, 2013; Halgren et al., 2002; Lau et al., 2013; 2014), the focus here was on the brain processes that support the explicit retrieval of strong associations (which are expected to be supported by their coherence with the structure of long-term semantic knowledge) and weak associations (which are less well-supported by long-term conceptual information and thus might require greater engagement of controlled retrieval processes to shape retrieval to suit the demands of the task). We examined how the retrieval of strong and weak semantic conceptual relationships was reflected in (i) changes in oscillatory power over time, as measured by MEG; and (ii) vulnerability to inhibitory online brain stimulation, using chronometric TMS.

In both experiments, the same behavioural paradigm was used to explore the functional and temporal organisation of semantic processing in the anterior and posterior temporal lobe (ATL and pMTG). Previous work has associated ATL with the retrieval of strong associations, in conjunction with other regions in the default mode network (Davey et al., 2016; Jackson et al., 2015), while controlled retrieval is thought to engage semantic control processes in pMTG, together with LIFG, to allow non-dominant aspects of meaning to come to the fore (Noonan et al., 2013; Badre et al., 2005; Gold et al., 2006; Davey et al., 2015; 2016). In line with these predictions, task-induced changes in oscillatory power were greater for strong than weak associations in ATL, while pMTG showed the opposite pattern (weak > strong associations). TMS confirmed a causal role for these sites in the efficient retrieval of strong and weak associations respectively. Timing differences between the sites were also found: ATL showed greater oscillatory power for the strong associations around 400ms post-target onset, although a strong task-related response was observed in the MEG data across conditions even before the onset of the second word (reflecting the successive presentation of multiple meaningful items in our paradigm). TMS to ATL disrupted performance for strong associations at around 150ms, around the time that early effects of semantic manipulations have been reported at this site in other studies (Clarke et al., 2011; 2012). This time point may have been sensitive to the disruptive effects of TMS (even though the difference between strong and weak conditions was not significant in the MEG data until later) since a coherent pattern of semantic retrieval was not yet fully established (and was therefore vulnerable to interference). pMTG showed an even earlier differential response to the strong and weak conditions in both MEG and TMS: this site responded more strongly to weak associations throughout the analysis window (from about 60ms post-onset of the second word), and TMS delivered to pMTG at the point of target onset impaired the efficient retrieval of weak associations. Thus, the MEG and TMS results followed the same temporal sequence across sites, although the critical time for TMS-induced disruption preceded the emergence of condition differences in MEG. Below, the contributions of ATL and pMTG to semantic cognition are discussed in light of these findings.

*Anterior temporal lobe*: The ATL is proposed to play a crucial role in heteromodal conceptual representation (alongside modality-specific ‘spokes’; Patterson, Nestor & Rogers, 2007; Rogers et al., 2006; Coutanche & Thompson-Schill, 2014). ATL is important for accessing conceptual knowledge from visual inputs (alongside other modalities) – a process that activates the ventral visual stream which terminates in ATL (Visser, Jefferies, Embleton & Lambon Ralph, 2012; Visser, Jefferies & Lambon Ralph, 2009). MEG studies of this aspect of ATL processing have identified responses in this region within 120ms of stimulus onset (Clarke et al., 2013; Fujimaki et al., 2009; Yvert et al., 2012). In addition, ATL is implicated in relatively automatic aspects of semantic access and retrieval (Lau et al., 2013; Davey et al., 2016). The current findings are highly consistent with this emerging story about the contribution of the ATL to semantic processing but add several important elements.

First, we used beamforming to characterise the response in ATL to strong and weak associations in total oscillatory power. In contrast, other MEG studies localising semantic effects to ATL have largely used measures maximally-sensitive to evoked power (Halgren et al., 2002; Bemis & Pylkkänen, 2011; Westerlund & Pylkkänen, 2014; Zhang & Pylkkänen, 2015; Lau et al., 2014; Fujimaki et al., 2009). Total power includes both phase-locked components and signals that are *not* phase-locked to the onset of the stimulus. Since the emergence of coherent semantic activation over time draws on long-term knowledge of the meanings of words across contexts, one might expect this process to generate neural oscillations that are not directly linked to stimulus onset. In line with these considerations, strong task-induced decreases in total power to the target were found in both ATL and pMTG. These effects were not seen in response to the presentation of the first word in the pair (see Figure 3), and therefore this response could be a marker of the retrieval of meaning at least partly decoupled from the stimulus itself. This interpretation draws on the view that power decreases are not necessarily associated with a decrease in neural activity (Hanslmayr et al., 2012; Hanslmayr, Staresina & Bowman, 2016): decreases in total power can reflect an increase in desynchronised neural activity that allows the representation of richer informational content, and our results can be interpreted within this framework – strong associations are more coherent with the structure of long-term conceptual knowledge and might generate richer or more meaningful experiences.

TMS to ATL disrupted the efficient retrieval of strong more than weak associations at 150ms post-stimulus onset – i.e., at the point when interactions between visual cortex and ATL are thought to become established (Clarke et al., 2011; 2012). In the MEG data, there was a strong task-related response in ATL by 150ms, although there was not yet a significant difference between the strong and weak conditions. Thus, the emergence of coherent semantic retrieval for the strongly-linked items may have been vulnerable to perturbation from TMS before the pattern of response within the ATL was well-established. Although a previous cTMS study found disruption when TMS pulses were applied to ATL at 400ms post-trial onset (Jackson et al., 2015), this study did not examine differential disruption of strong vs. weak associations, and it involved a more complex two-alternative-forced-choice decision as opposed to yes/no decisions about the presence or absence of a relationship between two words – thus the timings are unlikely to be comparable.

*Posterior middle temporal gyrus:* While the importance of ATL for conceptual representation is relatively widely accepted, there is considerable controversy about the role of pMTG in semantic cognition, since dominant theoretical frameworks have suggested that this site (i) represents particular aspects of lexical or semantic knowledge – such as event representations; or (ii) supports controlled semantic cognition as part of a large-scale network that includes LIFG. Studies have shown a common response in pMTG and LIFG using a wide range of manipulations of semantic control – including contrasts of ambiguous over non-ambiguous words, decisions with strong vs. weak distracters, and the retrieval of weaker versus stronger semantic links, in paradigms similar to the one adopted here (Noonan et al., 2013). pMTG is functionally connected to both the executive network and ATL, suggesting this region may be well-placed to control retrieval from the semantic store (Davey et al., 2016). Offline TMS studies have provided convergent evidence for the disruption of weak (but not strong) semantic association judgements when inhibitory stimulation is applied to pMTG (Whitney et al., 2011; Davey et al., 2015). When the relationship between probe and target is weak, the probe will tend to activate features and associations that are irrelevant to the decision that has to be made, and consequently semantic retrieval may have to be shaped to suit the task demands – irrelevant information must be suppressed while non-dominant aspects of knowledge are brought to the fore. The current data support the role of pMTG in controlled aspects of semantic retrieval: moreover, while previous studies have commonly focussed on showing *similarities* between LIFG and pMTG (refs: Noonan et al., 2013; Whitney et al., 2011a and 2011b), these findings show an important functional dissociation for the retrieval of weak and strong associations between anterior and posterior sites within the temporal lobe.

The time-course of these effects place important constraints on theories of controlled semantic retrieval: pMTG would be expected to show a relatively late response to the comparison of weak and strong if controlled retrieval takes time to become established, and if activity at this site reflects a re-interpretation or re-shaping of semantic activation following initial semantic retrieval driven by the written input. Alternatively, pMTG would be expected to show an early response to the same comparison if this site is important for maintaining information that is currently relevant and triggering the recruitment of the semantic control network when incoming information is not strongly coherent with ongoing semantic retrieval. pMTG may be able to reduce the propagation of dominant features and associations recovered from ATL when initial processing of new inputs suggests that these aspects of knowledge may be insufficient for comprehension. The current data support the second of these two alternatives. In MEG, the weak > strong effect commenced within 50ms of target-onset and continued throughout the analysis window. Using cTMS, evidence for an early role of pMTG was found in the efficient retrieval of weak associations, since there was greater disruption of weak trials when TMS was applied at target onset. The experimental design presented words sequentially (not concurrently), and consequently the findings are consistent with the hypothesis that pMTG maintains currently-relevant features or interpretations and detects situations in which incoming information is not well-aligned with these aspects of knowledge. This interpretation is consistent with studies that have shown a stronger response to more predictive primes in pMTG, including adjectives (Fruchter et al., 2015) and pictures (Dikker & Pylkkänen, 2013) that are informative about upcoming items. In our task, information about the semantic context might have been more critical for the efficient retrieval of weak associations, since it might have supported the rapid engagement of controlled retrieval processes when expectations were partially met. In contrast, for strong associations, relevant features in the semantic store will have been primed by the first word and thus this process may be less critical.

The visualisation of total power across the whole epoch in Figure 3 also revealed task-related decreases in power in pMTG that were marked by the offset of the prime and sustained throughout the interval between the two words. This observation is consistent with the interpretation offered above; that pMTG may maintain currently-relevant semantic information determined by the context, allowing controlled retrieval processes to be engaged at an early stage when inputs are not readily coherent with features that are being maintained. This perspective is further consistent with studies suggesting that pMTG shows strong engagement when meaningful inputs themselves determine a context that requires semantic retrieval to be shaped in a particular way (Davey et al., 2016; Badre et al., 2005).

Some limitations of this research are worth noting. First, this study focuses on the role of two key locations in the temporal lobe predicted to show a functional dissociation in the Controlled Semantic Cognition framework (ATL and pMTG). By combining targeted analysis of MEG data (examining local peaks within these regions) with chronometric TMS delivered to these sites, strong conclusions can be drawn about the nature of this dissociation, although the study is uninformative about other regions in the brain. Secondly, there is increasing evidence of functional subdivisions within ATL and pMTG (e.g., temporal pole, ventral ATL and aSTG appear to have different functional profiles – Lambon Ralph et al., 2017). The limited spatial resolution of MEG, and practical limits on the number of TMS sessions does not permit the separation of these regions. Thirdly, it may not be appropriate to directly compare timings across the MEG and TMS experiments, since Figure 1b demonstrates that the behavioural responses recorded within the MEG scanner were considerably slower than those obtained in the laboratory. This may have contributed to differences between our experiments; particularly the earlier effects of strength of association seen in the TMS study relative to the MEG study. More generally, this observation supports the view that it may not be possible to precisely specify the timing of neurocognitive responses, since these timings will critically depend on the task or paradigm that they are measured within. For example, the timing of differential responses to strong associations and weak associations might be influenced by experimental factors such as the stimulus-onset asynchrony (SOA), which is known to modulate the extent to which semantic priming draws on automatic or controlled processes (Gold et al., 2006). This study used brief stimuli presentation (200 ms) and a short SOA (150 ms), in order to limit the impact of factors such as stimulus repetition and proportion of related to unrelated trials (Neely, 1977; 1991). Furthermore, though the priming literature is relevant to our interpretations, our paradigm is not directly comparable to priming experiments, since we required participants to make an explicit judgement of the relationship between the two words, as opposed to examining the facilitatory influence of meaning on reading. An alternative approach, which we adopted here, is to consider the relative timing of behavioural effects *within a paradigm* which can then be localised to different brain regions.

Taken together, these results indicate dissociable roles of ATL and pMTG in semantic retrieval. ATL and pMTG showed opposite effects of strength of association in a semantic judgement task in both the MEG and cTMS experiments, supporting the proposal that these sites make a differential contribution to more automatic and controlled aspects of semantic retrieval.

## Acknowledgements

The research was funded by BBSRC (BB/J006963/1). EJ and NS were supported by the European Research Council (SEMBIND – 283530). JS was supported by the European Research Council (WANDERINGMINDS – 646927).

